# Benchmarking full-length transcript single cell mRNA sequencing protocols

**DOI:** 10.1101/2020.07.29.225201

**Authors:** Victoria Probst, Felix Pacheco, Finn Cilius Nielsen, Frederik Otzen Bagger

## Abstract

Single cell mRNA sequencing technologies have transformed our understanding of cellular heterogeneity and identity. For sensitive discovery or clinical marker estimation where high transcript capture per cell is needed only plate-based techniques currently offer sufficient resolution. Here, we present a performance evaluation of three different plate-based scRNA-seq protocols. Our evaluation is aimed towards applications requiring high gene detection sensitivity, reproducibility between samples, and minimum hands-on time, as is required, for example, in clinical use. We included two commercial kits, NEBNext*®* Single Cell/ Low Input RNA Library Prep Kit (NEB*®*), SMART-seq*®* HT kit (Takara*®*), and the non-commercial protocol Genome & Transcriptome sequencing (G&T). G&T delivered the highest detection of genes per single cell, at the absolute lowest price. Takara*®* kit presented similar high gene detection per single cell, and high reproducibility between sample, but at the absolute highest price. NEB*®* delivered a lower detection of genes but remain an alternative to more expensive commercial kits.

## Introduction

Within the last decade technologies for Single Cell Sequencing (SCS) has progressed research on tissue heterogeneity, cellular identity, and cellular state. Today, single cell technologies are applied at scale, famously in the Human Cell Atlas (HCA) project (Regev et al., 2017), and similar initiatives including Human Biomolecular Atlas Program (HuBMAP) from National Institute of Health (NIH) (HuBMAP, 2019), and The LifeTime Initiative (*https://lifetime-fetflagship.eu/*). Single cell mRNA sequencing (scRNA-seq) allows for the study of inter- and intra-cellular transcriptional variability, and delineation of transient cellular processes, identification cell types, marker genes and pathways. All current scRNA-seq techniques require isolation and lysis of single cells with subsequent conversion of RNA to cDNA and amplification of cDNA. Amplification is necessary due to the small amount of starting material, limited to mRNA content in a single cell and current, scRNA-seq protocols yields data that suffers from amplification bias (Ilicic et al., 2016). Library preparation for scRNA-seq vary a lot in terms of what information it’s possible to uncover from the data, and the protocol should be carefully chosen depending on the biological problem at hand (Ding et al., 2020; Ziegenhain et al., 2017).

The *modus operandi* for single cell sequencing is the addition of a unique tag (barcode) to the DNA/RNA from each single cell, which in turn allows for highly multiplexed sequencing on a short-read sequencer, like popular machines from Illumina. After sequencing demultiplexing allows for separation of data from each cell, using the barcodes. Techniques for scRNA-seq are roughly divided in two categories: plate-based and droplet-based techniques and starting material for both technologies is dissociated single cells in suspension (Fig. 1). In plate-based techniques, each single cell is plated alone into a chamber of a multi-well PCR plate (or into single tubes). Barcodes are added to each well, as a part of final library preparation and amplification. In droplet-based techniques, transcript barcoding takes place within the first step of the process, in a flow chamber making oil-emulsion droplets, with one cell per droplet. Recent commercial kits and flow chamber machines for droplet-based sequencing makes this protocol relatively easy entry but does not allow of sequencing of the full-length transcripts. Plate-based methods requires, in comparison, more technical know-how as well as separate handling of each cell. In practice plate-based methods only allows for processing of some hundreds of cells in parallel, whereas droplet-based methods can allow for preparation of thousands of cells in a single batch. Plate-based approaches are still more sensitive and allow for detection of more genes per cell and furthermore allows for additional protocols on same cell, such as quantification of FACS surface markers, and various DNA sequencing protocols (Moudgil et al., 2020; Svensson, Vento-Tormo, & Teichmann, 2018; Wang, He, Zhang, Ren, & Zhang, 2019) (Table 1).

**Table 1.** Table summarizing *pro et contra* of droplet versus plate-based techniques.

|  | Droplet-based | Plate-based |
| --- | --- | --- |
| Advantages | More Cells<br>Less hands on time<br>UMI-based | More Genes<br>Full-length cDNA<br>Enable collection months before processing |
| Disadvantages | Less Genes<br>3' - or 5'end cDNA<br>Require Immediate processing | Less Cells<br>More hands on time<br>No UMI's |

**Figure 1.**
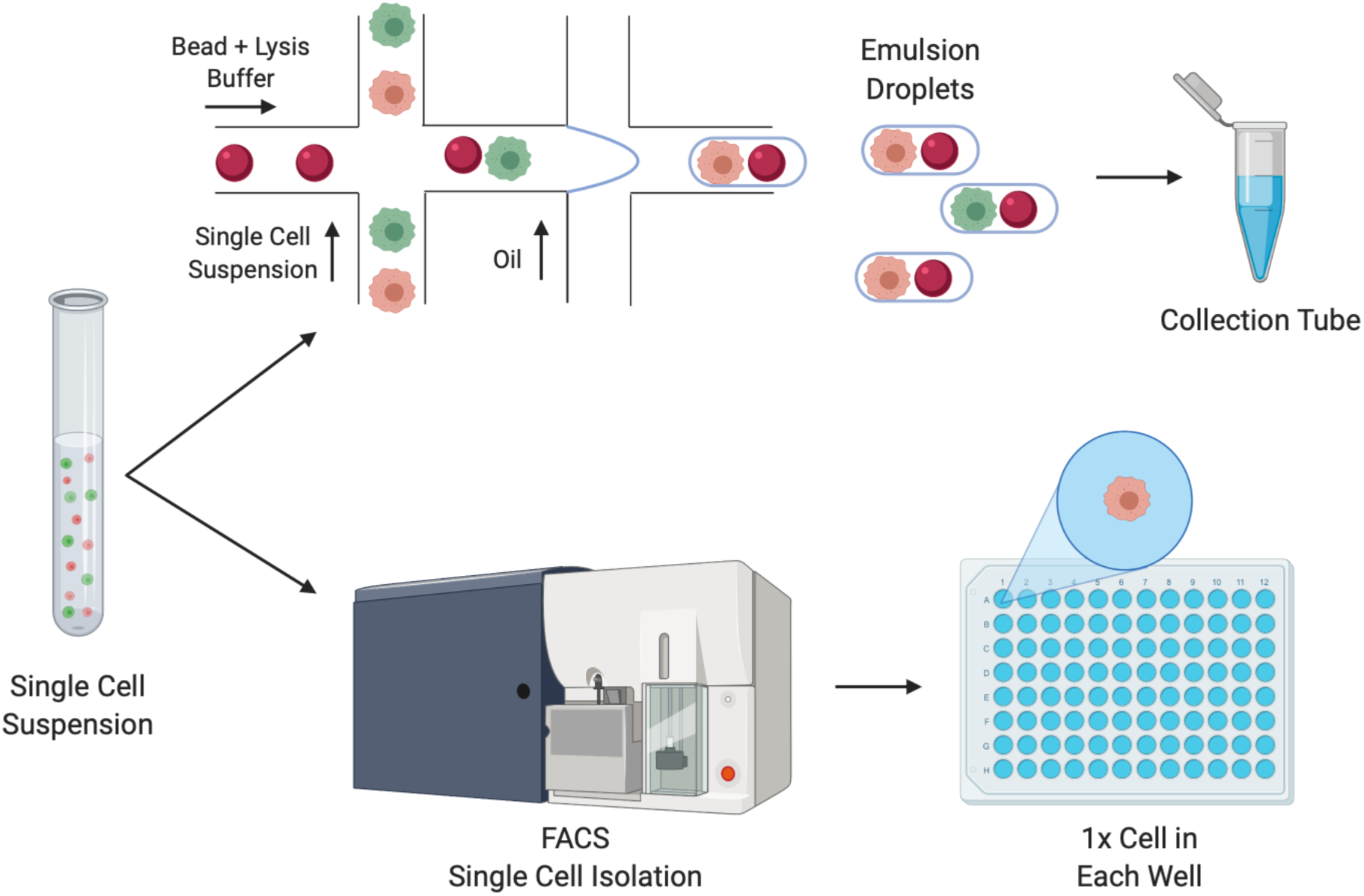
Workflow for droplet (top) and plate-based (bottom) single cell sequencing technologies. Droplet-based approaches combine primer-covered beads and single cells in emulsion droplets. Cells are lysed within the droplet, and reverse transcription carried out. Single cell cDNA is subsequently pooled and processed in bulk. Plate-based approaches isolate single cells into lysis buffer containing multi-well PCR plates or tubes. Isolation may be performed using fluorescent activated cell sorting (FACS). Barcodes are added during final library preparation steps.

In order to uncover structural variation, mutations within transcripts, detection of pseudogenes, and splice variants, sequencing of the full-length transcript is needed. Full-length transcript scRNA-seq techniques are currently all plate-based. A disadvantage of current full-length sequencing techniques is the preclusion of early barcoding and incorporation of Unique Molecular Identifiers (UMIs). Adding UMIs in an experiment aims to establish a unique identity of each RNA molecule (Islam et al., 2014). During PCR amplification, each cDNA containing the same UMI are assumed to be derived from the same mRNA molecule. Inclusion of UMI’s counting gives the protocol higher power with regards to transcript copy number detection (Fu, Wu, Beane, Zamore, & Weng, 2018; Grun, Kester, & van Oudenaarden, 2014).

SMART-seq (Switching Mechanism At the 5’ end of RNA Template) is a plate-based technique selectively capturing polyadenylated (poly(A)) RNA transcripts. The protocol yields libraries of full-length transcripts and relies on Reverse Transcription (RT) followed by template switching (TS) (Simone Picelli et al., 2014). In brief, the poly(A)-tail of mRNA transcripts are primed using an oligo-d(T) primer coupled to a PCR handle. The primed mRNA is reverse transcribed by Moloney Murine Leukemia Virus (M-MLV) RT, which has terminal transferase activity, and adds non-templated nucleotides to the 3’end of cDNA ends. These non-templated nucleotides are preferentially cytosines, which allow annealing of a template switching oligo (TSO) containing ribo-guanosines at its 3’end. The 2nd generation of SMART-seq, SMART-seq2, applies a TSO carrying a locked nucleic acid (LNA) at the 3’end. The LNA locks the nucleotide in endo-formation, which improves base-stacking and annealing ability and results in a higher melting temperature between the cDNA strand and the TSO (Grunweller & Hartmann, 2007). The LNA gives SMART-seq2 higher transcript capture, which results in improved sensitivity in gene detection (S. Picelli et al., 2013). SMART-seq2 has been reported to be the most sensitive and accurate method in terms of gene detection, and gives the most even read coverage, among all current scRNA-seq protocols (S. Picelli et al., 2013; Ziegenhain et al., 2017). Several different SMART-seq kits are commercially available differing in chemistry, price, and hands-on processing time. In this study, three different SMART-seq full-length protocols; NEBnext*®* Single Cell/Low Input RNA Library Prep Kit for Illumina (New England Biolabs (NEB*®*)), SMART-seq*®* High-Throughput (HT) kit (Takara Bio Inc.), and G&T-seq were performed on T47D cell line for comparison of sensitivity and precision between each protocol.

### NEBnext® Single Cell/Low Input RNA Library Prep Kit for Illumina

NEB*®* is a commercially available kit containing enzymes and buffers required to convert RNA, either purified or from cultured or primary cells, to cDNA for sequencing on Illumina platforms. The kit is plate-based (or tube) and builds upon the techniques of original SMART-seq. NEB ULTRA II FS DNA library preparation for preparation of Illumina sequencing compatible libraries is included with the kit. The protocol had a price tag of 46 € per single cell when processing 12 reactions (Table 2). Advantageously, this kit includes both Reverse transcription (RT), PCR amplification, and final library preparation with no further reagents needed for sequence-ready libraries compatible with sequencing on Illumina machines (Fig. 2 A). From here on, the protocol will be referred to as NEB*®*.

**Table 2.**
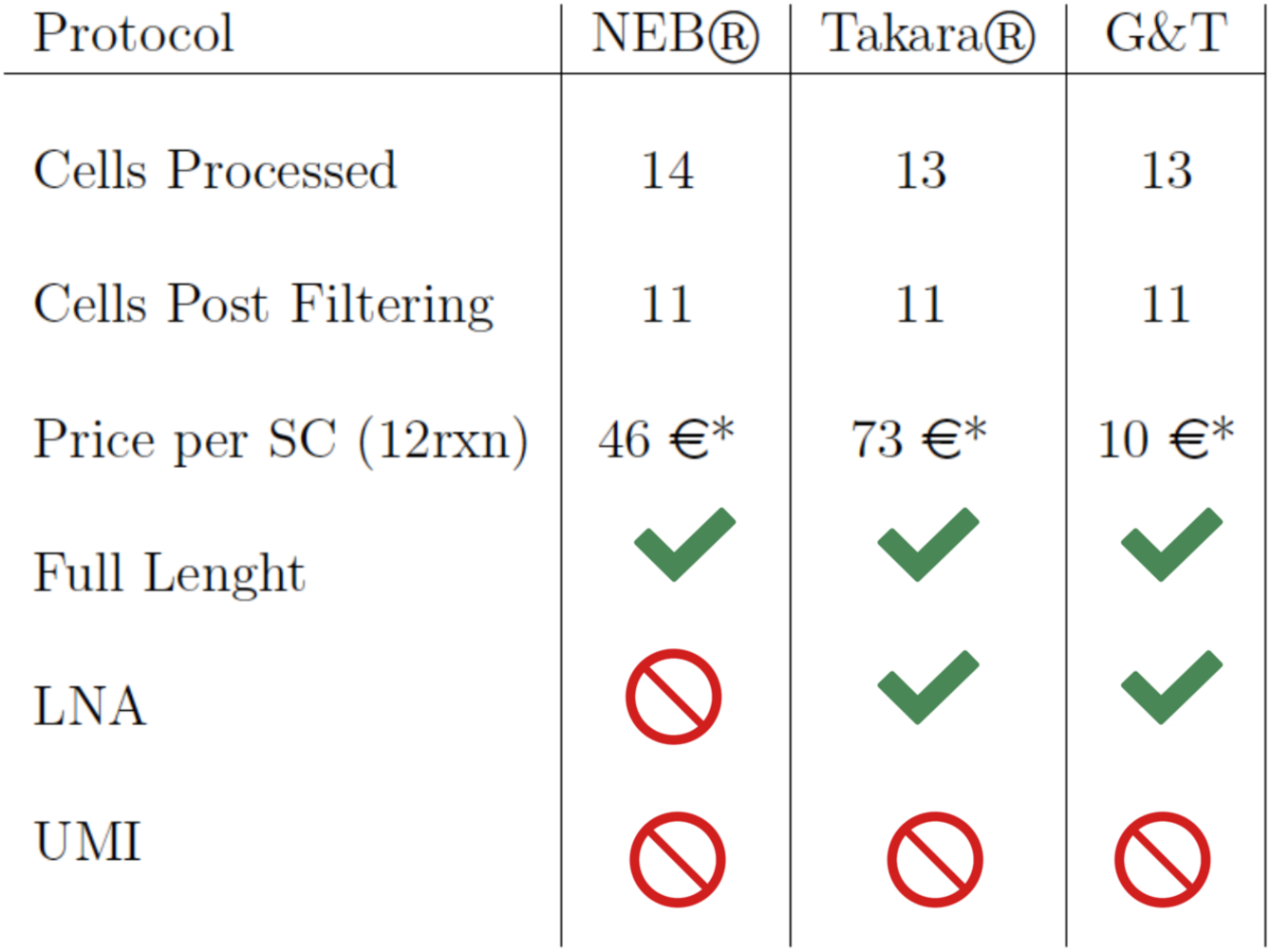
Table constituting the number of cells processed and cells remaining post QC filtering. All prices are per single cell using the smallest commercially available format of 12 cell reactions (rxn), as quoted medio 2020. All tested protocols feature full-length sequencing of transcripts. Takara® and G&T protocol genes a TSO with a locked Nucleic Acid (LNA), whereas NEB® doesn’t. None of the protocols feature UMI’s. *Price includes reagents for reverse transcription, PCR amplification, and final library preparation for all protocols. Price for protocols Takara® and G&T include price for final library preparation at 8 € per cell for 1/4 Nextera XT DNA library preparation (Cat. Nr: FC-131-1096) according to recommendation by kit manufacturer (*Takara Bio Inc*.) and protocol (G&T) (I. C. Macaulay et al., 2015).

**Figure 2.**
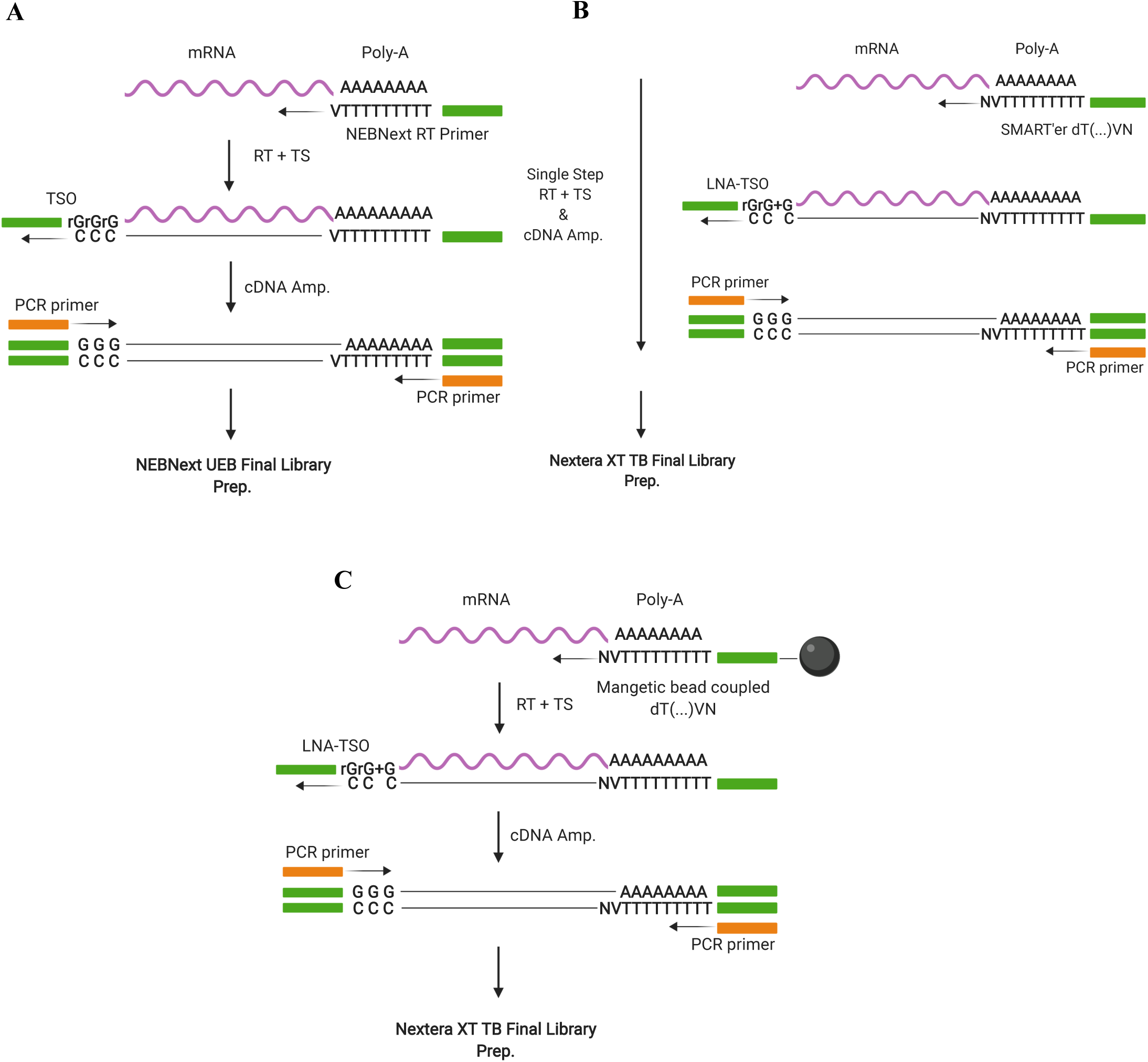
Illustration of the three scRNA-seq protocols applied in this study. A) NEBNext® Single Cell/Low Input RNA Library Prep Kit for Illumina (NEB®) followed by NEBNext® uracil excision based (UEB) Final Library Preparation KIT. B) SMART-seq® High-Throughput (HT) kit (Takara Bio Inc.) followed by final library preparation using Nextera XT Library preparation kit (Illumina, USA). C) Genome & transcriptome sequencing (G&T-seq) followed by final library preparation using Nextera XT Library preparation kit (Illumina, USA).

### SMART-seq® HT kit

SMART-seq*®* HT kit (Takara Bio Inc.) is commercially available and designed for generating full-length cDNA from single cells or purified total RNA. The mechanism of the reaction is built upon a patented version of SMART-seq. SMART-seq*®* HT applies the newer SMART’er technology, SMART-seq2. The kit differs from traditional SMART-seq2 protocol by combining RT and cDNA amplification in a single step, which minimizes hands-on time. In this comparison, final library preparation was performed using Nextera XT Library preparation kit (Illumina, USA). This protocol is the most expensive at 73 € per single cell when processing 12 reactions (Table 2). This price includes RT and PCR amplification, as well as final library preparation according to recommendations by manufacturer (Fig. 2 B). From here on, this protocol is referred to as Takara*®*.

### G&T Protocol

Genome & Transcriptome sequencing (G&T-seq) is not a commercially available protocol, and was originally developed for the study of both the genome and transcriptome of the same single cell (Iain C. Macaulay et al., 2016). G&T protocol has previously been shown to outperform traditional SMART-seq2 protocol (I. C. Macaulay et al., 2015; Svensson et al., 2017). The key divergence of the original SMART-seq2 method, is a step separating mRNA from genomic DNA (gDNA), into distinct single cell samples eligible for a range of library preparation protocols. This step also serves as an RNA purifying step, removing cell debris, protein and gDNA form the downstream reaction, and the method have been applied to hard-to-sequence cells, where the DNA fraction has been discarded (Choudry et al., 2020). The protocol originally applies a modified SMART-seq2 protocol for transcriptome amplification, and PicoPLEX or Multiple displacement amplification (MDA) for genome amplification (Iain C. Macaulay et al., 2016; Simone Picelli et al., 2014). The separation of gDNA from mRNA is enabled by an oligonucleotide containing a PCR sequence, a stretch of 30 thymidine residues (oligo-d(T)30), and an anchor sequence (VN)(V=A,G, or C; N=A,G,C or T) coupled to biotin in the 5’end. The 5’Biotin modification enables conjugation to streptavidin coated magnetic Dynabeads*®* (Fig. 2 C). Oligo-d(T)30VN beads capture mRNA transcripts in a cell lysate solution, and transcripts are moved to one part of the well using a magnet. This allows for transfer of gDNA in the solution to a new plate. Following separation, gDNA and mRNA is individually processed and sequenced, allowing for correlation of genomic mutations with gene expression. The G&T-seq workflow is not commercially available as a kit and is the cheapest (10 € per single cell), however also the most demanding protocol to set up (Table 2). Each reagent has to be individually purchased and solutions prepared. The RT step also requires specialized equipment (Eppendorf Thermomixer C) for on-bead SMART-seq2 conduction. Throughout this article the protocol is referred to as G&T.

### Quality Measures for single cell RNA-seq

Common quality metrices applied in scRNA-seq are library size and nr. of genes detected per cell. Library size is the total sum of mapped sequencing reads (counts) across all genes for a single cell. Library size mainly depends on sequencing depth, but given that an adequate number of reads have been obtained from sequencing, cells with small libraries can be considered of low quality. Small library size can be the result of RNA degradation due to either contamination (e.g. RNases), apoptosis, inefficient transcript capture before first strand synthesis or cDNA amplification. Importantly for our comparison, library size may also depend on protocol chemistry, reflecting the ability of adaptor-ligated fragments to anneal to oligos on the flow cell. Often of interest for scRNA-seq protocols is the ability to detect the vast repertoire of different gene transcripts within each single cell. The more genes a protocol is able to identify the more sensitive the protocol is evaluated. Here, we assess each protocol’s ability to detect endogenous mRNA transcripts of single cells.

To account for technical biases in a full-length RNA-seq protocol, External RNA Control Consortium spike-ins (ERCCs) can be added. ERCCs consist of 92 synthetic transcripts of bacterial origin that function to standardize sequencing experiments by adding an equal amount to each single cell reaction prior to processing steps. ERCCs show minimal sequence homology with endogenous eukaryotic transcripts, but features a poly-A tail, has different GC-content and vary in lengths (Jiang et al., 2011). The total proportion of reads mapping to ERCC spike-ins can be used to assess the quality of input cells, because it can be assumed to be inversely proportional to good-quality fragments from the cell which are available for sequencing (Ilicic et al., 2016). Applying ERCCs in a sequencing experiment can also be used to account for biases such as primer capture-efficiency, batch effects and absolute RNA content estimation, because the same amount and concentration is added to each sample. The proportion of mitochondrial (MT) mapping reads, can also be used as a QC metric for cell quality (Ilicic et al., 2016), the reason being that transcripts within mitochondria are better protected both from leakage and degradation (Ilicic et al., 2016; Islam et al., 2014). The different mitochondrial genome consist of 37 genes, and captured MT genes can be assessed as an endogenous capture-efficiency control the same way synthetic ERCCs spike-ins are used.

## Results

### Data Generation

Data was obtained by sequencing cells from breast cancer cell line T47D. mRNA from 13 single cells was amplified using Takara*®* SMART-seq*®* High-Throughput (HT) kit and G&T-seq protocol, while mRNA from 14 single cells was processed using NEB*®* NEBNext*®* Single Cell/Low Input RNA Library Prep Kit for Illumina (Table 2). For retrieval of homogenous cell populations, cells were stained for, and selected by, expression of human EpCAM and Integrin *α*6 (CD49f) (Fig. 3 A, supplementary Fig. 1) (Gligorich et al., 2013; Keller et al., 2010). Cells were sorted by FACS into lysis buffer compatible with each protocol (Methods). Cells had same cell line origin and yielded similar results on measured parameters, and major variability between cells was for all protocol cell cycle stage (Fig. 3 A, supplementary Fig. 2). We therefore consider each cell of same identity and use cells as biological replicates for benchmarking robust sequencing results. Single cells were sequenced in groups according to processing protocol on Miseq (Illumina, USA), aiming at a sequencing depth of one million reads for sufficient capture of genes across protocols (Fig. 3 B). Following quality control (QC) and filtering, 11 cells from each protocol were kept in each dataset (Table 2). Processing of Cells and sequencing was performed uniformly for all tested protocols.

**Figure 3.**
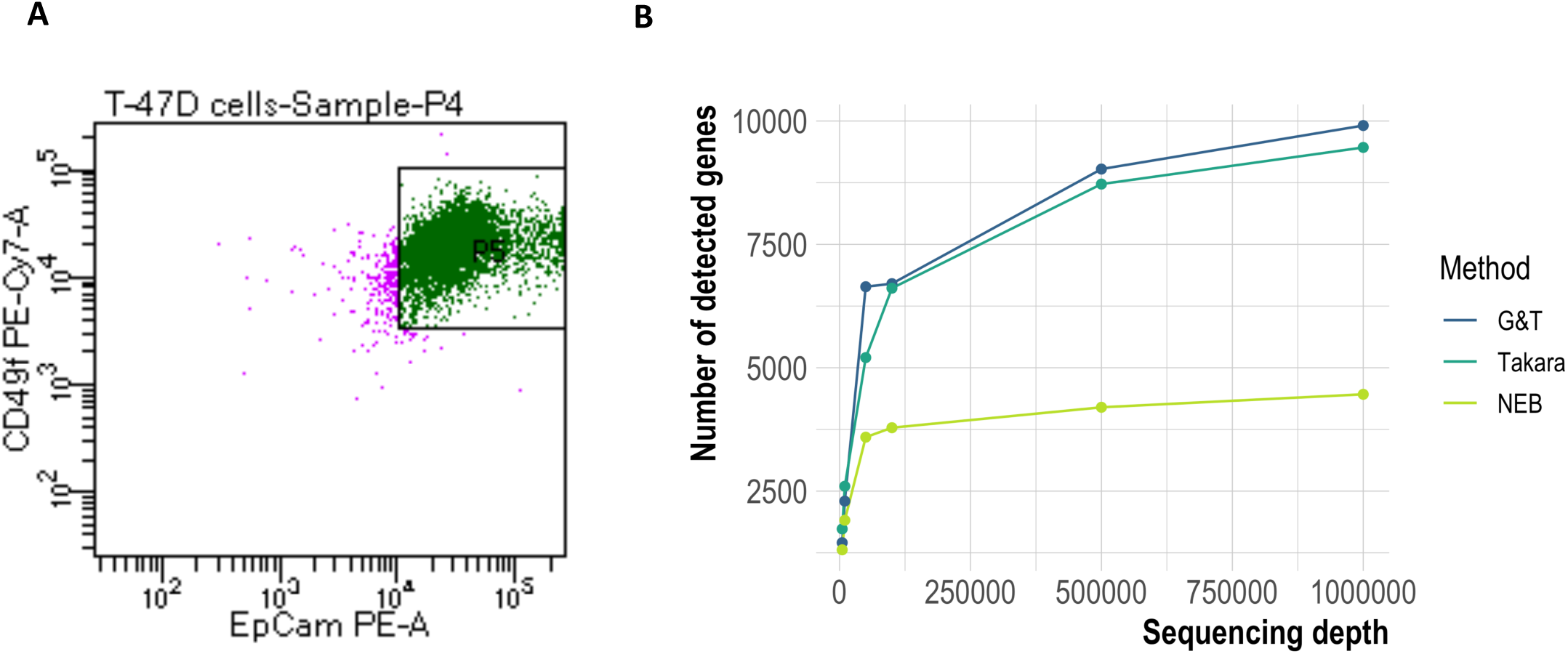
Cell Selection & Gene Saturation. A) Homogenous EpCAM+/CD49f+ T47D cells sorted into lysis buffer containing 96-well plates by fluorescent activated cell sorting. B) Gene capture saturation prior to filtering performed by resampling subsets of total reads to 5000, 10.000, 50.000, 100.000, 500.000 and 1 million reads per cell using seqtk (https://github.com/lh3/seqtk).

### Takara® had highest cDNA yield, NEB® had lowest

In order to assess the RT and PCR amplification step, the total amount of cDNA was measured from each single cell library post RT and PCR amplification. Takara*®* and G&T processed cells were subjected to 20 cycles of PCR amplification, whereas 22 cycles were necessary for successful amplification of NEB*®* processed cells. Cells processed using Takara*®*, and G&T had a total cDNA yield of ∼ 84ng (*σ* = 19 ng) and ∼ 59 ng (*σ* = 7 ng) per single cell, respectively. (Fig. 4 A). NEB*®* cells had the lowest average total cDNA yield of ∼ 39 ng per single cell (*σ* = 20 ng) (Fig. 4 A), despite the increased number of PCR cycles applied.

**Figure 4.**
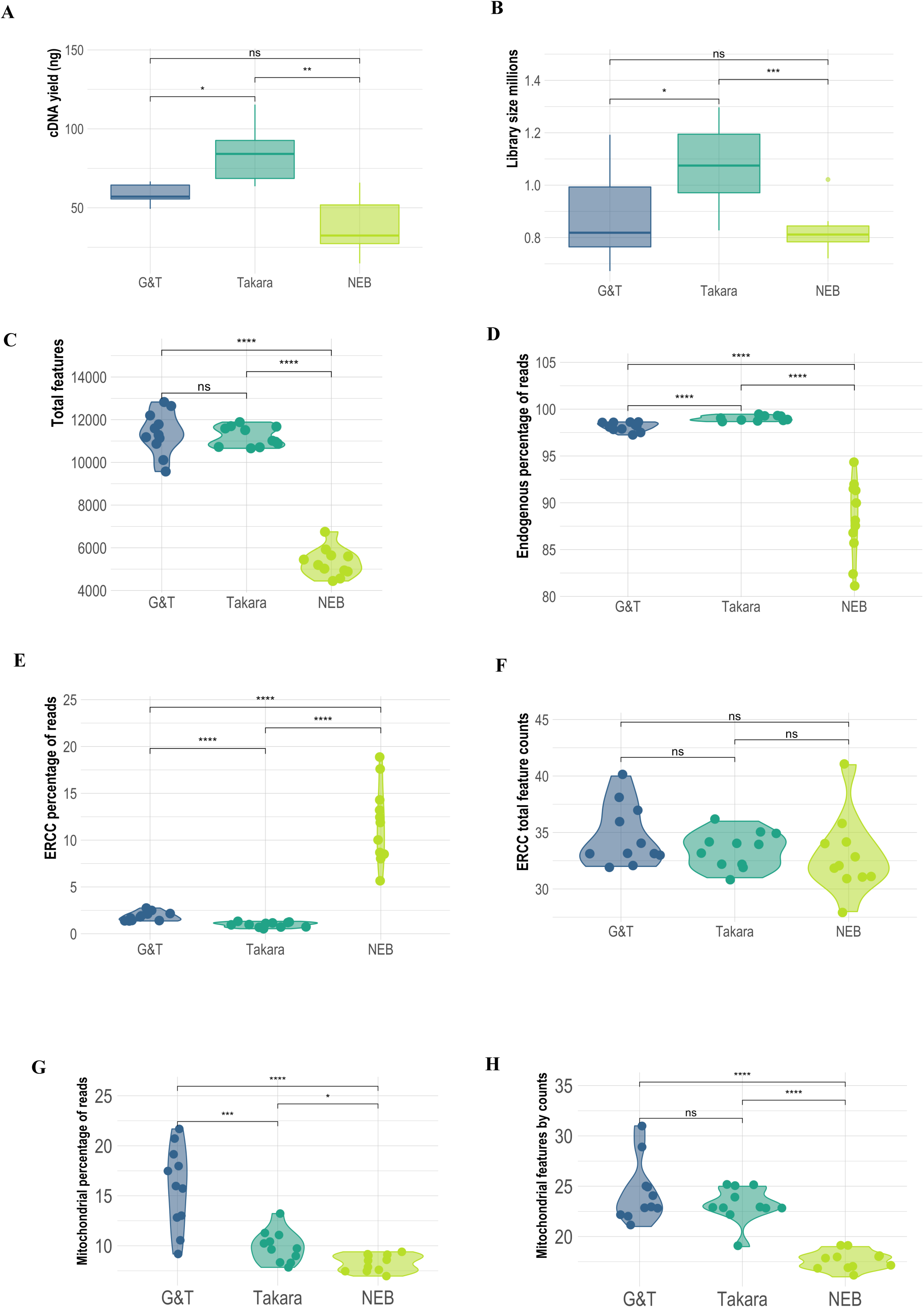
Comparative quality metrices from scRNA-seq data generated with NEB®, Takara®, and G&T protocols. A) cDNA ng/yield following mRNA amplification in each protocol. B) Library sizes millions. C) Total nr. genes detected per single cell. D) Percentage of reads mapping to endogenous genes per single cell. E) Percentage of reads mapping to exogenous genes; ERCC spike ins per single cell. F) Nr. Captured ERCC genes per single cell. G) Percentage of reads mapped to mitochondrial (mt) genes per single cell. H) Nr. Captured MT genes per singe cell. Each dot represents a single cell.

### Takara® cells had largest libraries, G&T and NEB® had similar library sizes

In order to evaluate amplification efficiency we assessed library size of each protocol. Takara*®* cells had significantly (p < 0.02) larger libraries than remaining protocols with an average library size of 1.08 × 10^6^ reads per cell (*σ* = 0.2 × 10^6^ reads) (Fig. 4 B, Table 3). G&T and NEB*®* cells had an average library size of 0.88 × 10^6^ reads per cell (*σ* = 0.2 × 10^6^ reads) and 0.82 × 10^6^ reads per cell (*σ* = 0.8 × 10^5^ reads), respectively (Fig. 4 B, Table 3). However, NEB cells showed the most consistent library sizes yield across cells, between 0.72 – 1.02 × 10^6^ (Fig. 4 B, Table 3).

**Table 3.** Table constituting summary of average library size, total genes captured, capture-efficiency of control genes; MT and ERCC’s per single cell following filtering for protocols NEB®, Takara®, and G&T.

| Protocol | NEB® | Takara® | G&T |
| --- | --- | --- | --- |
| Lib. Size (Millions) | 0.82 | 1.06 | 0.88 |
| Total Features | 5.310 | 11.211 | 11.382 |
| MT Features | 18 | 23 | 24 |
| ERCC Features | 33 | 34 | 35 |

### Takara® & G&T had highest capture-efficiency of control genes, NEB® had the lowest

Low fraction of reads mapping to ERCC and MT genes is a marker of high quality and robustness in single cell protocols (Ilicic et al., 2016). Takara*®* cells had the lowest average proportion of reads mapped to ERCC spike-ins at 0.97% (*σ* = 0.26%), with G&T and NEB*®* 1.86% and 11.75%, respectively. The ability to recapture ERCC was similar across protocols with an average of 35 by both G&T and Takara*®* and 33 for NEB*®*, out of the 92 added spike-ins. (Fig. 4 E F, Table 3). G&T, Takara*®*, and NEB cells had an average proportion of reads mapped to MT transcripts at 15.84% (*σ* = 1.6%), 9.91% (*σ* = 1.6%), and 8.3% (*σ* = 0.84%), respectively. Average MT transcripts covered per cell for G&T, Takara*®*, and NEB was 24 (*σ* =3,07), 23 (*σ* =1.72), and 18 (*σ* =0.92) transcripts per cell out of 37 MT genes. (Fig. 4 G H, Table 3). The correlation between MT and ERCC varied among protocols spanning from positive correlation (G&T: 0.9, Takara*®*: ρ = 0.33) to negative correlation (NEB*®*: ρ = −0.64).

### Takara® & G&T cells had the highest average gene count per single cell, NEB® cells had the lowest

In order to evaluate gene capture efficiency, the total number of genes with at least one read mapped was assessed in each single cell. Takara*®*, G&T and NEB produced libraries with an average of 99% (*σ* = 0.26%), 98% (*σ* = 0.47%), and 88% (*σ* = 4.09%) genes mapped to endogenous genes (Fig. 4 D). Cells processed by G&T-seq recaptured the highest average number of 11.382 uniquely expressed genes per single cell (*σ* = 990) compared to Takara*®* Kit (11.211, *σ* = 463), and NEB*®* (5.310 genes, *σ* = 661), with only half the coverage (Fig. 4 C, Table 3).

### Data from Takara® cells were most consistent over cells, while NEB® had highest variance

In order to evaluate reproducibility between samples in each protocol, similarity between single cells was assessed by Pearson correlation coefficient (PCC) between all cells from the same protocol. NEB*®* processed cells were least similar with an average PCC of ρ = 0.58 (*σ* =0.02) Takara*®* and G&T cells showed significantly higher similarity between cells with an average PCC value of ρ = 0.86 (*σ* = 0.02) and ρ = 0.80 (*σ* = 0.03) (p << 0.0001) (Fig. 5 A).

**Figure 5.**
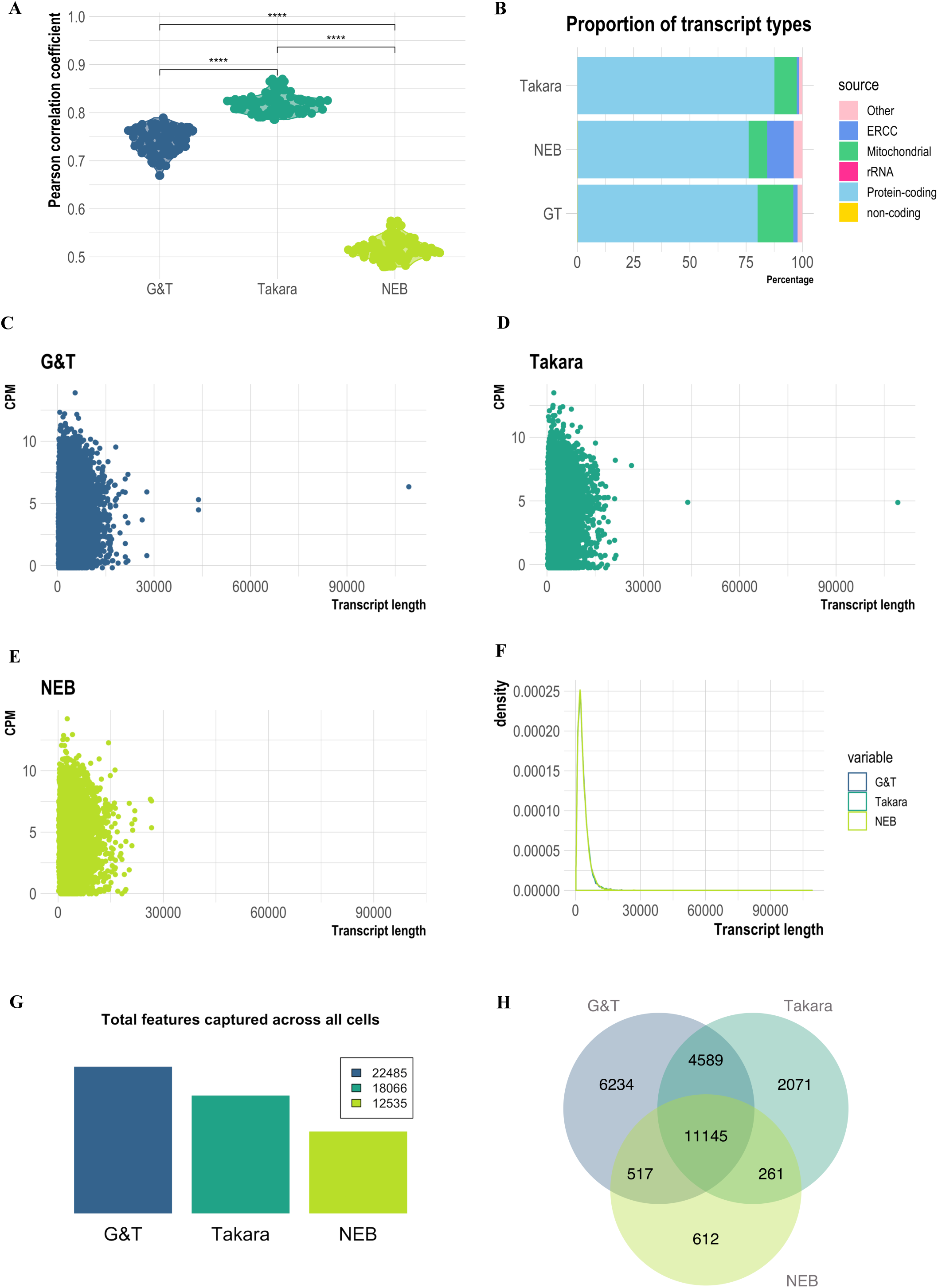
Comparative analysis of cell similarity, transcript type, length and unique genes between scRNA-seq data generated with NEB®, Takara®, and G&T protocols. A) cDNA ng/yield following mRNA amplification in each protocol.. B) Proportion of different transcript types in each library. Non-coding transcript types constitute e.g. tRNA, snRNA, snoRNA, miRNA, miscRNA, lincRNA. Distribution of detected transcripts by gene length (bp); C) G&T, D) Takara, E) NEB®. F) Density of transcripts according to length (bp) across all tested protocols. G) Bar-chart illustrating total unique genes captured across all cells in all three protocols. H) Venn diagram visualizing overlap of total captured genes between protocols, and genes captured uniquely in each protocol.

**Figure 6.**
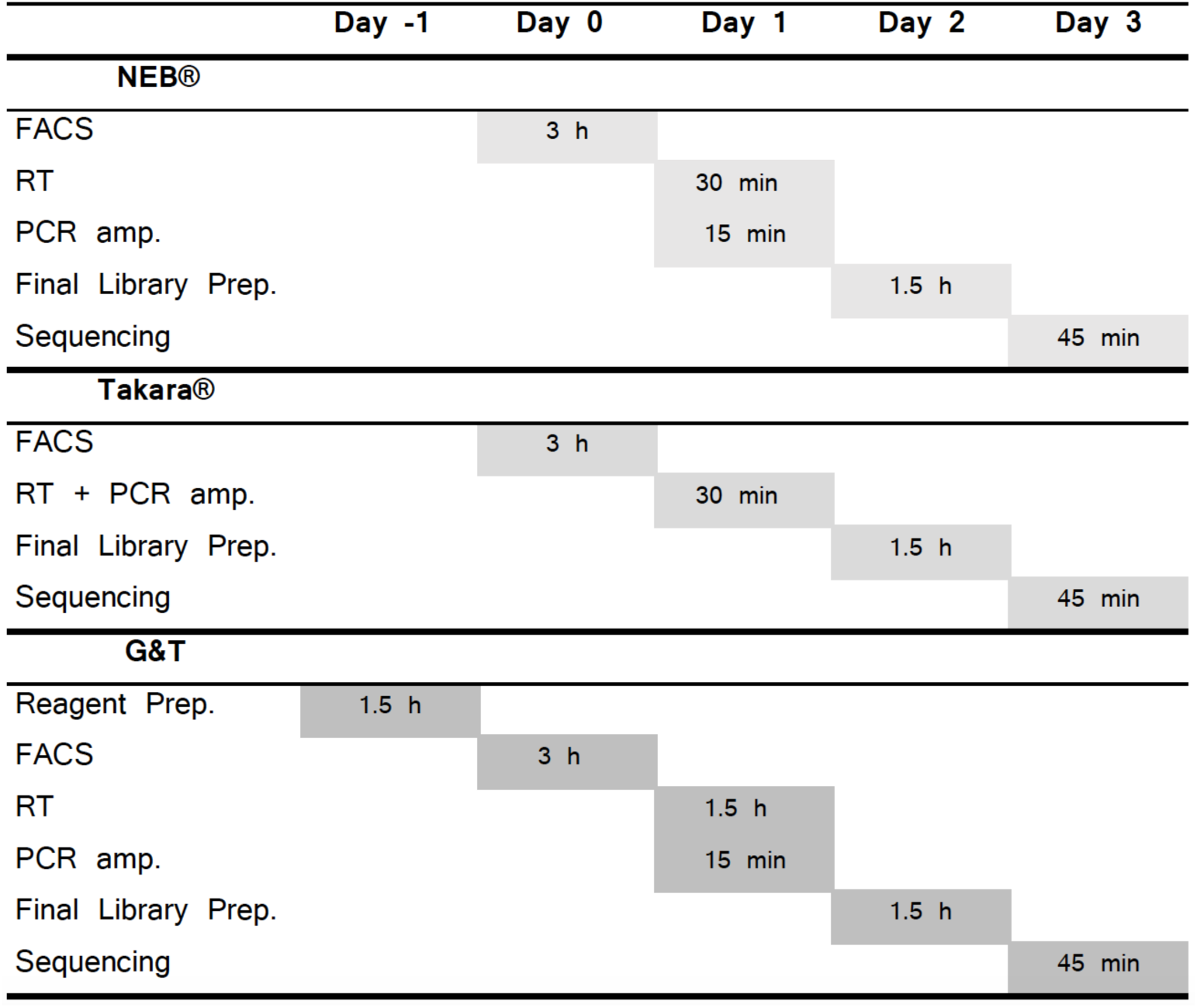
Hands-on time per protocol. Gantt chart featuring estimated hands-on time of each step of each tested protocol. NEB® protocol has approximately 6 hours of hands-on time. Takara® protocol has approximately 5 h and 45 minutes of hands-on time, by combining RT and PCR amp. Experimentalist save time both on processing but also leaves option for leaving the plate in thermocycler overnight, potentially saving even more time. G&T protocol has the longest processing time at 8 h 30 min due predominantly to reagent preparation and the separation step between RNA and DNA. All times are estimates assuming a skilled experimentalist with no prior experience with the protocol.

### G&T cells captured the greatest number of unique genes across all cells

In order to evaluate the ability of each protocol to capture the vast repertoire of different transcripts, we assessed proportion of transcripts belonging to different biological categories, the total amount of genes captured by each protocol, as well as the amount of genes captured uniquely in each protocol (Fig. 5 B G H). Proportion of captured transcripts was similar between all tested protocols (Fig. 5 B). However, Takara*®* had the largest proportion of transcripts mapping to protein-coding genes with an average of 88% (*σ* = 1.8%) compared to 80% (*σ* = 4 %), and 76% (*σ* = 3.6%) for G&T and NEB*®*, respectively (Fig. 5 B). G&T samples had the highest amount total genes captured across all cells at 22.485 genes (Takara*®*: 18.066, NEB*®*: 12.535) (Fig. 5 G). Additionally, G&T cells had the largest number of genes which were uniquely captured by that protocol with 6.234 genes across all cells (Takara*®*: 2.071, NEB*®*: 612) (Fig. 5 H). Finally, neither of the tested protocols had a transcript length bias, demonstrating that mRNA molecular quantification was not influenced on this measure in either of the protocols (Fig. 5 C-F).

## Discussion/Conclusion

This study is a performance evaluation of three different plate-based scRNA-seq protocols; NEBNext*®* Single Cell/ Low Input RNA Library Prep Kit (NEB*®*), SMART-seq*®* HT kit (Takara*®*), and Genome & transcriptome sequencing (G&T) (Fig. 2). G&T protocol was found most sensitive in regards of gene detection, with the highest detection of genes per single cell (avg. 11.382 genes per cell) but not significantly different from Takara*®* processed cells (avg. 11.211 genes per cell) (p = 0.48) (Fig. 4 C, Table 3). Furthermore, G&T captured the greatest number of genes across all single cells (22.284 genes), as well as having most genes uniquely captured by this protocol (6.234 genes) (Fig. 5 G H). This suggest that G&T protocol might be superior in detecting rare transcripts across cells for improved unravelling of heterogeneity within a complex population of cells (e.g. tumor tissue). Takara*®* protocol showed a high gene detection similar to G&T protocol (avg. 11.211 genes per cell) (Fig. 4 C, Table 3). Takara*®* and G&T were found most consistent (PCC = 0.86 and PCC = 0.80), suggesting high degree of reproducibility, especially important in e.g. a clinical setting (Fig. 5 A). NEB*®* protocol was found least sensitive in regard to gene detection, both per single cell (avg.5.310 genes per cell) and across all cells (12.535 genes) (Fig. 4 C, Table 3). Low gene detection might lead to false positive detection of heterogeneity within a cell population, induced by undetected genes and not by true biology. In certain studies, as well as in diagnostics, quantification of single marker genes can be of great relevance, thus a high degree of undetected genes cannot be tolerated.

High correlation between ERCC and mtRNA reads for G&T (ρ =0.9) would suggests robust quality metrics, whereas negative correlation in the case of NEB*®*(ρ = −0.64) could be a result of a competitive situation either during capture or sequencing. However, since the high performing Takara kit display little correlation (ρ =0.33) between ERCC and mtRNA, we speculate that several quality metrices are needed to give a full picture of quality, as also suggested by *Ilicic et al. 2016*. Thus, results are likely affected by more than just competition for capture or reads. The technical differences between these protocols are believed caused partly by chemistry (e.g. LNA vs. no-LNA, lysis buffer, reverse transcriptase etc.) and reaction volume (lower volume = higher sensitivity) (Svensson et al., 2017). None of these protocols addresses PCR amplification biases, however, not yet commercially available technology SMART-seq3 is the newest generation protocol for full-length scRNA-seq, and implements UMI’s in the 5’end of full-length RNA transcripts (Michael Hagemann-Jensen, 2019). Inclusion of UMI counts gives the protocol higher power in regard to transcript copy-nr. detection (Michael Hagemann-Jensen, 2019). Furthermore, SMART-seq3 has been suggested to improve the sensitivity of original SMART-seq protocols to levels approaching single-molecule RNA fluorescence in situ hybridization (smRNA FISH) (Michael Hagemann-Jensen, 2019). Plate-based Quartz-seq2 and microfluidic 10xchromium 3’end RNA-seq are two examples of UMI featuring technologies that have previously performed well in regards to gene detection at low read depth (Mereu, 2020). However, the maximum nr. of captured genes remain at best one fourth lower for Quartz-seq2 and one fith lower for 10xchromium 3’end RNA-seq, compared to G&T and Takara*®* protocols featured in this study (Montoro et al., 2018; Sasagawa et al., 2018; Shnayder et al., 2018). 10xchromium 3’end RNA-seq technology, has also previously shown severe transcript drop-out risk, especially of rare transcripts (Wang et al., 2019). Furthermore, both Quartz-seq2 and 10xchromium are not full-length protocols, limiting detection of analysis across all exons, fusion-transcripts, splice-variants, as well as SNP mutation analysis which are especially relevant in e.g. studies of disease (Brinkman, 2004; Di Gregorio et al., 2013; Zhao, Hoadley, Parker, & Perou, 2016). 10xchromium 3’end RNA-seq allows parallel sequencing of up to ∼ 80.000 cells in a single run, whereas plate-based methods are limited most often to 96-well or 384-well format. Choosing protocol may therefore often boil down to the choice between high number of processed cells vs high number of detected genes.

Takara*®*’s SMART-seq*®* HT kit was evaluated as having the greatest ease-of-use, due to low hands-on-time, achieved by combining the step of reverse transcription and PCR amplification (Table 6). The ampure cDNA clean-up step of this protocol was also less time consuming and required fewer steps than NEB*®*’s Single Cell/ Low Input RNA Library Prep Kit. However, Takara*®*’s kit featured the highest price per single cell (73 € Per cell/ sample), and did not include reagents for final library preparation on Illumina machines. NEB*®*’s kit included reagents for RT, PCR amplification, and final library preparation on Illumina machines. Even though NEB*®* did not outperform either G&T or Takara*®* on single cell metrices, chances are this kit may perform well as a low input RNA-seq protocol - as a cheaper alternative to Takara*®*’s kit (46 € per cell/ sample). G&T protocol is not commercially available, and it is the cheapest tested protocol (10 € per cell). G&T is also the most technically challenging to set up and require some specialized equipment; the separation step of G&T-seq may be performed manually using a magnetic plate, but for more high throughput experiments, experimenter may wish to use a programmable liquid handling robot. The RT step of G&T-seq furthermore requires a Thermomixer C (Eppendorf, cat. no. 5382 000.015), to prevent bead precipitation during on-bead RT reaction.

Overall, our comparison found that G&T processed cells showed the highest sensitivity in gene detection and high reproducibility at the absolute lowest price. However, G&T was the most time-consuming and most technically challenging protocol. Takara*®* processed cells showed a likewise high sensitivity in gene detection similar to G&T processed cells, however at the absolute highest price. Takara*®* protocol had the greatest ease-of-use, lowest hands-on-time, as well as highest reproducibility across single cells. NEB*®* cells showed lowest sensitivity of gene detection, and lowest degree of reproducibility between single cells. However, NEB*®* protocol had the advantage of including reagents for both RT, PCR amplification and final library preparation. In conclusion, we would recommend anyone with the skills and patience to perform G&T-seq due to high sensitivity at a low price. If you are new to the field Takara*®* offers a lower entry level protocol with high gene detection and high reproducibility across single cells, however at a higher price.

## Materials & Methods Single

### Cell Suspension

T47D single cell suspension was prepared by removal of growth medium and subsequent washing of cell layer using 10 ml PBS. Cells were disaggregated with 2 ml TrypLE™ Express Enzyme (1X) (cat nr. 12604013, Gibco™, USA) for approximately 2 min in an incubator at 37 C°. Reaction was stopped adding full growth media (RPMI-1640+GlutaMAX™ (cat nr. 61870-010, Gibco™, UK) + 10% Fetal Bovine Serum (FBS) + 5% Penicillin/ Streptamycin) in double the amount of TrypLE™. Suspension was spun down 3 min 1200 rpm. Cells were washed once in PBS, resuspended in FACS buffer (PBS + 0.04 % bovine serum albumin (BSA)), and filtered using a 100 μm strainer.

### Cell Surface Marker Staining

Cell suspension was stained with monoclonal antibodies toward human EpCAM (cat nr. 130-111-116, lot nr. 5190125111, Miltenyi Biotec, Germany) conjugated to R-phycoerythrin (PE) (Ex-Max 496 nm/ Em-Max 579 nm), and Human Integrin *α*6 (CD49f) (cat nr. 25-0495-80, lot nr. 4319156, eBioscience™, USA) conjugated to PE-cyanine 7 (PE/Cy7, Ex-Max 496 nm/ Em-Max 785 nm). 4’,6-diamidino-2-phenylindole (DAPI) (Ex-Max 358 nm/ Em-Max 461 nm)(cat nr. 10236276001, Roche Diagnostics GmbH, Germany) was used to discriminate dead from live cells. Concentration was adjusted to 10^5^-10^6^ cells/ml, and antibody added at concentration of 1 μl per 10^5^-10^6^ cells. Cells were incubated 15 min at 4 C° in the dark. Following incubation, dye was diluted, adding 1 ml of FACS buffer. Suspension was subsequently washed once with 1 ml FACS buffer. Pellet was resuspended in 1 ml FACS buffer. 10μl DAPI was added per 1 ml cell suspension, to a final concentration of 300 nM, 5 min prior to sorting.

### Single Cell Sorting

Fluorescence activated cell sorting (FACS) was used for isolation of single T47D cells into lysis buffer containing 96-well semi-skirted PCR plates using instrument BD FACSAria*™* III (BD bioscience, USA). Controls for calibrating instrument included unstained cells, cells stained with DAPI, and beads compensating for spectral overlap between fluorochromes PE and PE-CY7, using MACS Comp Bead Kits Anti-REA (cat nr. 130-104-693, lot nr. 5181012289, Miltenyi Biotec, Germany) and Anti-RAT (cat nr. 130-107-755, lot nr. 5181015526, Miltenyi Biotec, Germany). Cell suspension was gated to isolate double positive (EpCAM^+^/CD49f^+^) single T47D cells. A Multi-cell positive control (50 cells) and an empty-well negative control (0 cells), and at least one RNA-control diluted to 10pg/ul were provided per plate. Following sorting cells were thoroughly vortexed, spinned down 1 min, flash-frozen in dry-ice and subsequently stored at −80 C°.

### NEBNext® Single Cell/ Low Input RNA Library Prep Kit

14 single cells were processed using NEBNext*®* Single Cell/ Low Input RNA Library Prep Kit (cat nr. E6420S, New England Biolabs (NEB), USA), and were sorted into 5 μl NEBNext Cell Lysis buffer (0.5 μl NEBNext Cell Lysis Buffer (10x), 0.25 μl Murine RNase Inhibitor, 4.25 μl H2O). Protocol was performed according to recommendation by manufacturer with minor changes - each single cell lysate were added 1 μl 1:10^6^ dilution of ERCC spike ins (cat nr. 4456740, Invitrogen, Thermo Fischer Scientific, Lithuania) prior to RT and PCR amplification was performed applying 22 cycles.

### SMART-seq® HT Kit

13 single cells were processed using SMART-seq*®* HT kit (cat nr. 634862, Takara Bio Inc, USA), and were sorted into 12.5μl FACS dispensing solution (0.95 μl 10xLysis buffer, 0.05 μl RNase Inhibitor,1 μl 3’SMART-Seq CDS Primer II A, 10.5 μl Nuclease-free H2O). Protocol was performed according to recommendation by manufacturer with minor changes - each single cell lysate were added 1 μl 1:10^6^ dilution of ERCC spike ins (cat nr. 4456740, Invitrogen, Thermo Fischer Scientific, Lithuania) prior to RT, and PCR amplification was performed applying 20 cycles.

### G&T-seq

13 single cells were processed by G&T-seq and were sorted into 2.5 μl RLT Plus buffer (cat nr. 1053393, Qiagen, Germany). ScRNA-seq was performed as described by *Macaulay et al*., *2016*. Single cell DNA sequnecing featued in this protocol was not conducted in this experiment, but DNA stored at −80 C°. Each single cell lysate were added 1 μl 1:10^6^ dilution of ERCC spike ins (cat nr. 4456740, Invitrogen, Thermo Fischer Scientific, Lithuania) prior to RT. PCR amplification was performed applying 20 cycles.

### Sequencing

T47D single cell cDNA libraries were sequenced in groups according to each library preparation protocol (13 or 14 single cells per run) on MiSeq Benchtop Sequencer (Illumina, USA), using MiSeq Reagent kit v2 300 cycles (cat nr. MS-102-2002, Illumina, USA). Prior to sequencing each single cell library was diluted to a concentration of 4 nM in EB buffer + 0.1% Tween 20. Prior to sequencing 3 μl of each 4 nM library was pooled in an Eppendorf tube. 5 μl 4 nM pool was mixed with 5 μl 0.2 nM NaOH and incubated 5 min at RT, for denaturing of double stranded cDNA. The denatured sample pool was diluted to a concentration of 20 pM by mixing 10 μl 2nM sample pool with 990 cold Hybridization Buffer 1 (HT1). Finally, 20 pM sample pool was diluted to a concentration of 10 pM, by mixing 500 μl 20 pM sample pool with 500 μl cold HT1.

### Alignment/ Trimming

Illumina sequencing raw reads were exported as fastq files. Fastq files were processed on a bash shell. Two rounds of trimming was performed using TrimGalore!. First trimming removed Nextera XT adaptors (”CTGTCTCTTATACACATCT”), and second trimming removed cDNA amplification adaptors (”AAGCAGTGGTATCAACGCAGAGT”). Quality assessment of sequencing output was performed by two rounds of FastQC after each trim. Trimmed sequences were aligned using pseudoaligner ‘Spliced Transcripts Alignment to a Reference’ (STAR) to Genome Reference Consortium Human Build 38 (GRCh38). STAR output files comprising reads per gene for each sample, which were merged in a single tab delimited file.

### Data visualization

Introductory figures Illustrated using *Biorender* (https://biorender.com/). ScRNA-seq data was imported into R studies as an expression matrix. The count matrix was transformed into a Single Cell Experiment Object (SCE-object) using Rstudios (v3.6.1 Opensource, https://rstudio.com/) package *SingleCellExperiment* (v1.6.0). Data graphs were generated using *ggplot2* (v3.3.0). Plots are either Sinaplots (Sidiropoulos et al., 2018) or box-plots where with outlier threshold at X and wiskers at Y, stars are denoted based on Wilcoxon test p-value; ns: p > 0.05, *: p < = 0.05, **: p < 0.01, ***: p < 0.001, ****: p < 0.0001.

### Single Cell Quality Control (QC)

Expression level of genes were quantified by CPM (counts per million). Genes with an average expression above zero (CPM > 0) across all cells were kept in the dataset. Cells not expressed in any cell (CPM = 0) were filtered away. Bad quality cells (or empty wells) were filtered away based on the following criteria: 1) cells that had less than 1000 uniquely expressed genes, 2) cells that had library sizes below 0.6e^6^ million reads, 3) cells that had more than 30% reads mapped to mitochondrial genes, and 4) cells that had more than 25% of reads mapped to ERCC spike-inn genes. Data analysis was performed with RStudio (version 3.6.1) using Bioconductor [https://www.bioconductor.org] packages (SingleCellExperiment, scater (McCarthy, Campbell, Lun, & Wills, 2017), sinaplot (Sidiropoulos et al., 2018), ggplot2, GenomicFeatures (Lawrence et al., 2013), sincell (Julia, Telenti, & Rausell, 2015), TxDb.Hsapiens.UCSC.hg19.knownGene, SummarizedExperiment, robCompositions, splatter (Zappia, Phipson, & Oshlack, 2017), reshape2, ggforce, gdata, hrbrthemes, viridis, VennDiagram, DESeq2 (Love, Huber, & Anders, 2014), dplyr, tidyverse (Wickham et al., 2019), gtable, gridExtra, hrbrthemes, ggforce, ggpubr) following guidelines from [https://scrnaseq-course.cog.sanger.ac.uk/website/index.html].

## Author Contributions

VP and FOB conceived and designed the study. VP, FP and FOB, analysed the data. VP performed the experiments. VP and FOB wrote the manuscript. FOB and FN supervised the studies. All authors read and approved the final manuscript.

## Data availability

Data analyzed in this paper is publicly available from GEO accession number GSE155506.

## Supplementary figures

**Supplementary Figure 1.**
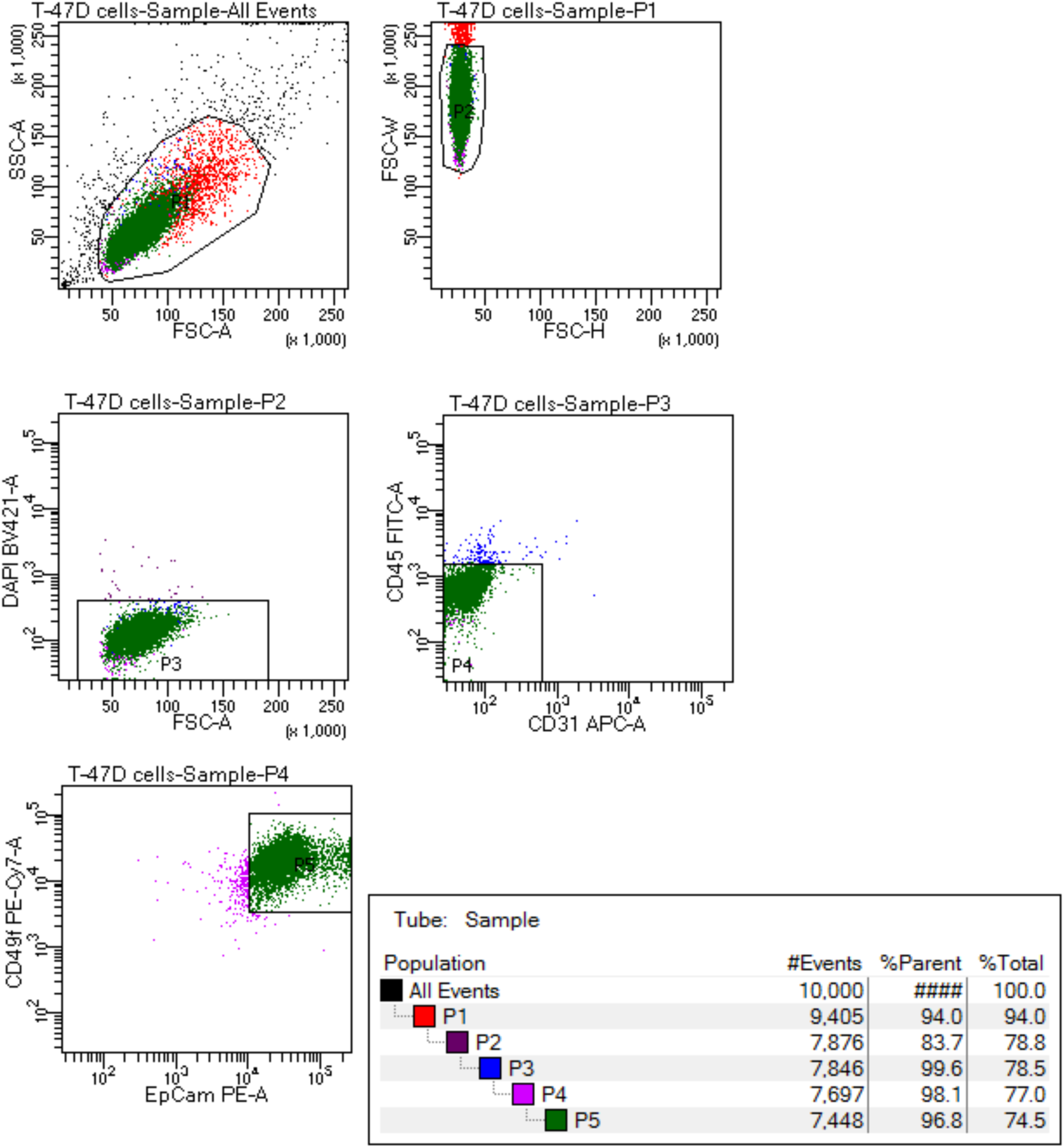
Fluorescent activated cell sorting (FACS) of single T47D cells. Cells were stained for EPCAM-PE, CD49f-PE/Cy7, CD31-APC, CD45-FITCH. P5: Double positive EPCAM+/CD49f+, sorted into lysis buffer containing 96-well plates according to protocol. P1: Population of live single T47D cells. P2: Singlet T47D cells, discarding doublets. P3: DAPI negative T47D cells, discarding dead/dying cells. P4: CD45-/CD31 negative population, control of non-specific binding of antibodies towards immune and endothelial cells.

**Supplementary Figure 2.**
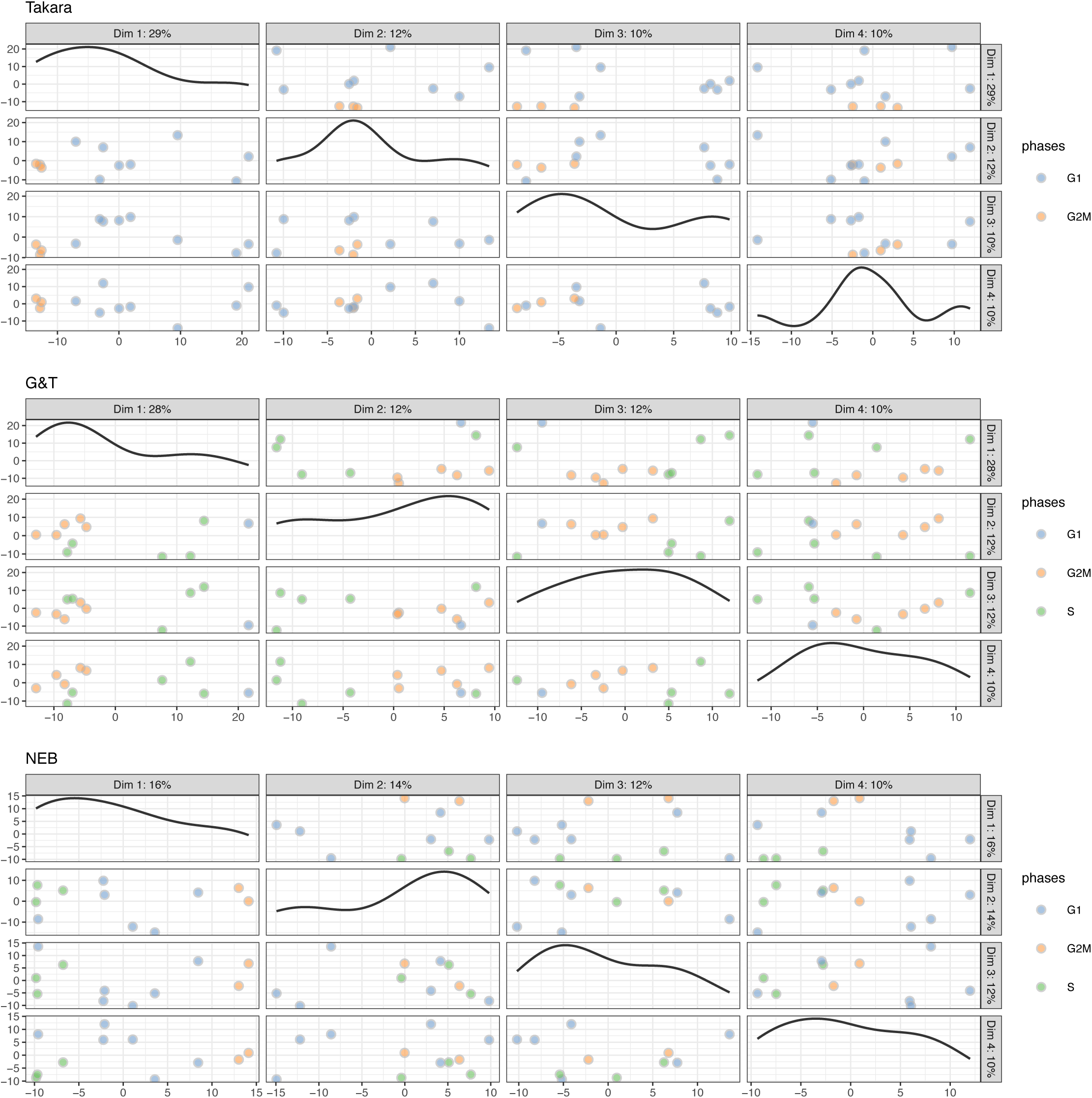
Cell cycle distribution between single cells of protocol Takara®, G&T and NEB®. Cell cycle assignment performed using cyclone (Scialdone et al., 2015).

